# Cortico-ocular coupling in the service of episodic memory formation

**DOI:** 10.1101/2022.11.09.515902

**Authors:** Tzvetan Popov, Tobias Staudigl

## Abstract

Encoding of visual information is a necessary requirement for most types of episodic memories. In search for a neural signature of memory formation, amplitude modulation of neural activity has been repeatedly shown to correlate with and suggested to be functionally involved in successful memory encoding. We here report a complementary view on why and how brain activity relates to memory, indicating a functional role of cortico-ocular interactions for episodic memory formation. Recording simultaneous magnetoencephalography and eye tracking in 35 human participants, we demonstrate that gaze variability and amplitude modulations of alpha/beta oscillations (10-20 Hz) in visual cortex covary and predict subsequent memory performance between and within participants. Amplitude variation during pre-stimulus baseline was associated with gaze direction variability, echoing the co-variation observed during scene encoding. We conclude that encoding of visual information engages unison coupling between oculomotor and visual areas in the service of memory formation.

## Introduction

Encoding of visual material is a prerequisite for most of our episodic memories and typically begins with the exploration of the environment. Relying heavily on vision, humans tend to explore the environment by eye movements. These typically include, but are not limited to, micro- and macro saccades as well as fixations jointly contributing to the overall gaze pattern^1^. Evidence suggests a strong association between gaze patterns and episodic memory formation^2-7^. Visual scenes that were explored with more eye movements are more likely to be remembered than scenes explored with less eye movements^8^.These associations between memory encoding and gaze are complemented by studies showing that gaze pattern reinstatement during retrieval supports successful recollection^9-12^. Similarly, gaze patterns are reinstated during visual imagery ^12-15^ even without the guidance of retinal input, i.e. during full darkness^16^.

Electro- and magnetoencephalographic studies confirm a robust power modulation of alpha/beta oscillatory activity (appr. range 10-20Hz) during the encoding of items that is predictive of later memory performance: alpha/beta power during later remembered items is reduced as compared to later forgotten items, an observation termed “subsequent memory effect” (SME^17^). Alpha/beta SMEs have been replicated consistently^18-23^, with remarkable specificity of alpha/beta power decreases during successful memory encoding^24-30^. This is of particular relevance as these alpha/beta effects have been interpreted as the mechanism by which the cortex tracks and organizes representations of the stimulus input to be fed forward to downstream memory circuits. According to these ideas, alpha/beta activity is functionally involved in successful episodic memory formation^24,31,32^.

Complementing such cognitive interpretations, a recent account of alpha/beta activity argues for a view closer to the biological implementation of the association between alpha/beta power modulations and oculomotor action^33^. Building on recent evidence demonstrating this close relationship in a variety of tasks^34-36^, the account postulates a fundamental and domain-general functional organization linking visual cortical signals to oculomotor action. The topographic modulation of alpha power (i.e. power reduction) was associated with consistent eye movements towards the contralateral visual hemifield^36-38^. We here set out to test the link between eye movements and alpha/beta activity in the context of episodic memory formation.

We hypothesize that alpha/beta power modulations link to episodic memory formation through consistent biases in gaze patterns. Thirty-five participants performed a memory task while simultaneously tracking eye movements and monitoring brain activity using magnetoencephalography (MEG). Gaze bias analyses revealed a robust gaze-related SME confirming that higher levels of visual exploration are associated with better memory performance. Crucially, high subjective confidence in correctly memorizing an item was predictive of gaze-related and alpha/beta SME. Moreover, splitting trials according to high versus low gaze bias revealed the typical alpha/beta SME over posterior sensors. During the baseline interval prior to scene encoding, spontaneous fluctuations of alpha/beta power reliably predicted the gaze direction variability, akin to the one observed during scene encoding. Exploratory analyses of the covariation between gaze direction and alpha/beta power confirmed a consistent relationship present throughout the entire recording and evident between and within participants.

## Results

### Memory performance evident in both gaze variability and power modulation of visual cortex activity

During the study phase, participants viewed visual scenes presented for 4 seconds. For each trial and participant, the simultaneously recorded eye tracking data were extracted and converted into 2D density heat maps with the x-axis representing the horizontal, y-axis vertical visual eccentricity. The density of gaze direction locations is color coded. Subsequently, a statistical contrast using non-parametric testing with clusters was performed^39^. This analysis revealed a robust gaze-related SME (Figure 1A, cluster-permutation test, p < 0.025; corrected for multiple comparisons across time and frequencies), indicating that, during the study phase, items that will be later remembered are characterized by a strong gaze bias away from fixation and towards various locations of the viewed scene. Quantification of the different eye movement types such as saccades and fixations using the EEG-EYE toolbox^40,41^ yielded similar results (supplemental Figure S1). For later remembered items, participants made more fixations (M/STD = 2044/544) and saccades (M/STD = 2070/545), as compared to later forgotten items (fixations: M/STD= 691/350; saccades: M/STD = 703/364). A significant condition difference was confirmed for both, fixations (t^34^ = 9.7, CI[1068 1637], p = 2.6977e-11, Cohen’s d = 1.64) and saccades (t^34^ = 9.7, CI[1079 1653], p = 2.7457e-11, Cohen’s d = 1.63). In the test phase, participants correctly recognized 72.83 % (+/-2.51 SEM) of the scenes on average, yielding a d-prime of 2.24 (+/-0.09 SEM).

**Figure 1:**
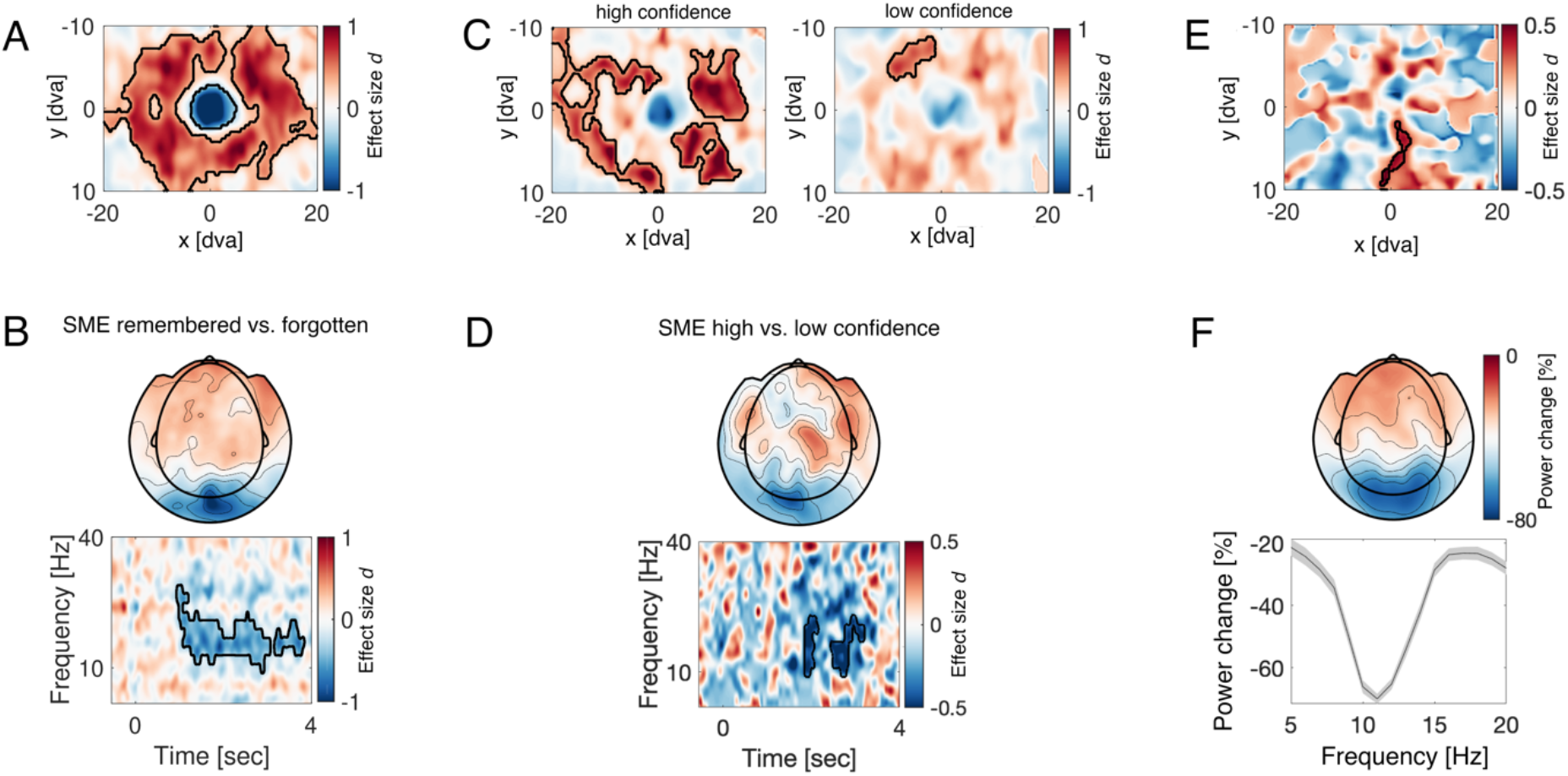
Coinciding subsequent memory effects (SME) reflected in gaze biases and modulation of visual alpha/beta power: **A-** Gaze density contrast between later remembered vs. later forgotten items. The 2D density maps were calculated per trial and latency between 0-4sec following scene presentation during the study phase. Red color indicates increased biased towards the respective visual screen locations for later remembered items and blue color decreased gaze density or gaze direction away from the respective location. Color code is expressed in units of effect size Cohen’s d obtained after cluster-permutation test (p< 0.025). **B**-Scalp topography and time-frequency representation of power (TFR) of the contrast between later remembered vs. later forgotten items during the study phase. Color code indicates effect size Cohen’s d obtained after cluster-permutation approach at p < 0.025. The black outline in the TFR highlights the cluster supporting the rejection of the null hypothesis of no effect between later remembered and later forgotten items. **C-** Similar to A but split according to the particpants confidence as of how certain their response is for an item being “old” or “new”. Gaze bias for high confidence trials (left) versus gaze bias for low confidence trials (right). **D**-2×2 Interaction SME (remembered, forgotten) x Confidence (high, low) illustrating the scalp topography and the TFR expressed in units of effect size Cohen’s d (cluster-permutation test, p < 0.025). **E-** Contrast in gaze density between trials dominated by low alpha/beta power minus high alpha/beta power during the prestimulus baseline (-1 to 0 sec) in the study phase. Color code is expressed in units of effect size Cohen’s d obtained after cluster-permutation test (p< 0.025). Red color indicates increased bias towards the respective visual screen locations for trials dominated by low alpha/power power as compared to high alpha/beta power. **F-** Scalp topography of the relative alpha/betapower change between the same trials as in (E) dominated by low vs. high alpha/betapower. Color code denotes the relative change in %. Statistical contrast is inapproriate in this case as it will be highly signficant per construction. This is however not the case for the evaluation of the gaze biases illustrated in E. The power spectrum of the difference averaged across occipital sensors is provided with shading denoting SEM.

Applying a standard (i.e. in accordance with previous work in the field) subsequent memory analyses on the MEG data confirmed the well documented alpha/beta SME (Figure 1B). Encoding of later remembered material was associated with a stronger alpha/beta decrease over occipital sensors (cluster-permutation test, p < 0.025; corrected for multiple comparisons across time and frequencies). The co-variation of stronger scene exploration (i.e., stronger gaze biases away from fixation for later remembered items) and modulation of visual alpha/beta power (i.e., stronger decrease for later remembered items) was informative for the participant’s memory confidence. Items reported with high confidence in being correctly memorized displayed stronger gaze biases (exploration) during encoding (Figure 1 C left, cluster-permutation test, p < 0.025) and alpha/beta SMEs (Figure 1 D, cluster-permutation test, p < 0.025), as compared to items with lower confidence (Figure 1C right). A potential alternative interpretation of the apparent relationship between eye movements and the modulation of visual alpha/beta power might be the subjective saliency of the stimulus material. Salience is hard to quantify objectively and strongly depends on subjective experience. Hence, some stimulus material could trigger both, variability in eye movements and alpha/beta power modulations. It could allow the conjecture that the apparent correlation between the two is driven by external stimulus factors. In order to test this alternative, we set out to explore a potential relationship between gaze variability and power modulation of ongoing alpha/beta activity using the data of the pre-stimulus (e.g. baseline) intervals during the study phase, where only a fixation cross instead of a visual scene was displayed. Participants were required and were successful in maintenance of central fixation, ranging within ±5° of horizontal and vertical visual angle (see Figure S2). We reasoned that, within participants, one could split the trials into high and low alpha/beta power (e.g. based on median split) during the pre-stimulus baseline interval. Subsequently, one could evaluate the distribution of gaze density around the required fixation. If the above conjecture is true, there should be no difference in the gaze variability between high and low alpha/beta power conditions, since the stimuli in the baseline period (that is, the fixation cross) did not vary in saliency. Conversely, differences in gaze bias would further confirm a co-varying relationship between the direction of gaze and modulation of visual alpha/beta power, independent of the task context and stimulus properties. The results of this analysis are illustrated in Figure 1 E and F. Within participants, trials with less alpha/beta power were also associated with a gaze pattern that deviated more from fixation (Figure 1 E). Conversely, high alpha/beta power during baseline was associated with reduction of the degrees of freedom in eye movements, resulting in less variability of gaze around the targeted (fixation) direction. This result echoes the observations made during stimulus presentation (e.g. Figure 1 A,B), both in terms of the direction of the relationship (i.e. alpha/beta power decrease relates to higher gaze variability) and scalp topography (Figure 1F).

### Gaze variability predicts both modulation of visual cortical activity and memory performance

Next we asked to what extend the inverse direction of high vs. low gaze variability will predict alpha/beta SMEs. Within participants we median split the trials during encoding into high and low gaze variability within the visual display area of high variability identified in Figure 1A (i.e. positive cluster). Trivially, the group difference in gaze variability is significant per design (Figure 2A). Importantly however this should not necessary hold for cortical alpha/beta power and/or memory performance. We observed that trials dominated with high gaze variability were also associated with stronger modulation of alpha/beta power over occipital sensors, akin to the alpha/beta SME topography observed in Figure 1B (cluster-permutation test, p < 0.02). Moreover, high gaze variability during encoding was associated with an increase in the number of later remembered items as compared to low gaze variability (Figure 2C, left). The opposite relationship was observed for forgotten items (Figure 2C, right). An alternative illustration of the same result is depicted in Figure 2D. The effect size of the memory contrast (# items remembered vs. forgotten) was dependent upon the gaze exploration during encoding: high gaze variability is linked to better memory performance as compared to low gaze variability (Figure 2D).

**Figure 2:**
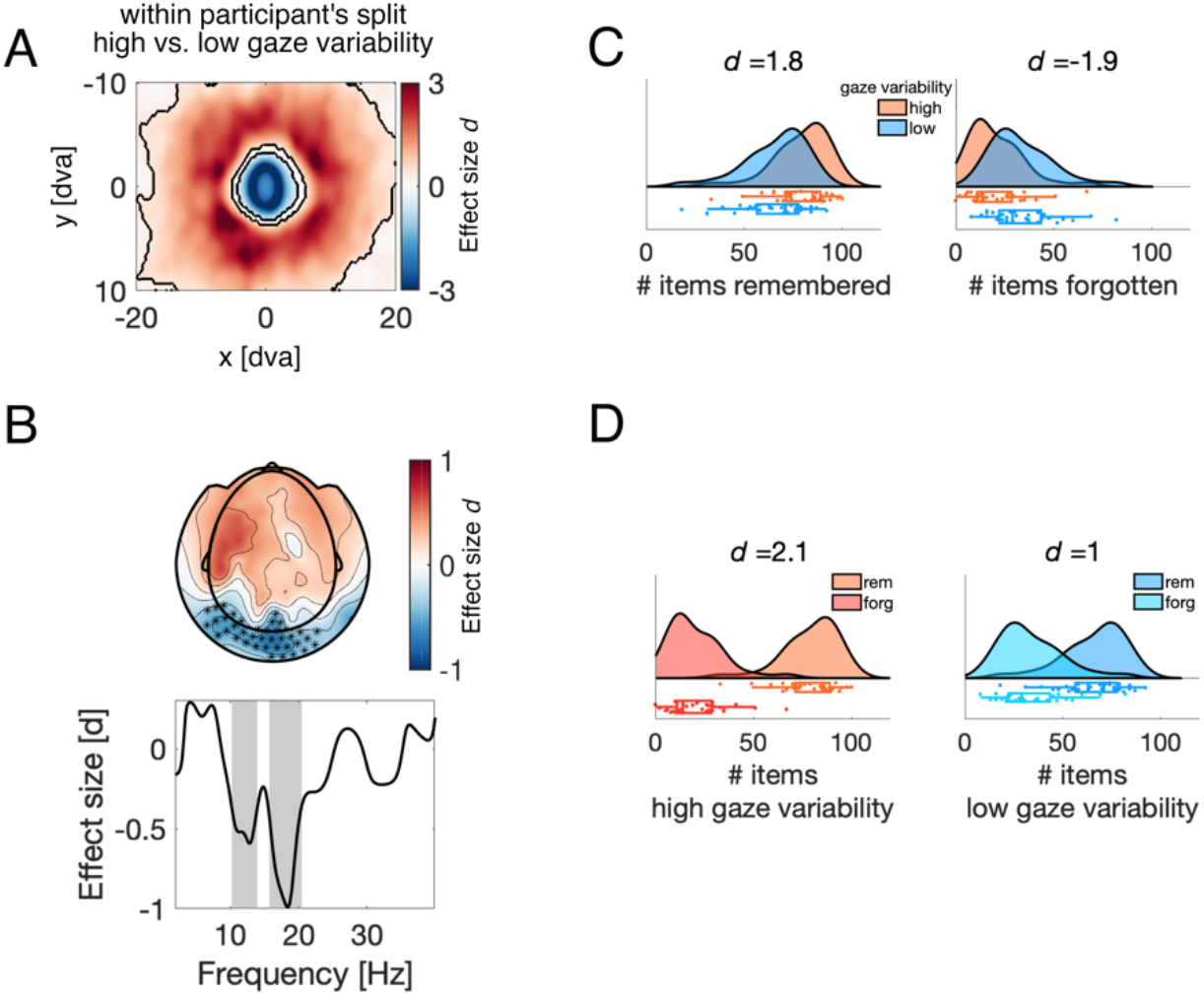
Gaze variability affects cortical alpha/beta power modulation and the effect size of memory performance. **A**-Gaze density contrast between trials dominated by high vs. low gaze variability (median split within participants). The 2D density maps were calculated per trial and latency between 0-4sec following scene presentation during the study phase. Red color indicates increased biased towards the respective visual screen locations for later remembered items and blue color decreased gaze density or gaze direction away from the respective location. Color code is expressed in units of effect size Cohen’s d obtained after cluster-permutation test (p< 0.025). **B-** Scalp topography and power spectrum contrast between high vs. low gaze variability trials during the study phase. Color code indicates effect size Cohen’s d obtained after cluster-permutation approach at p < 0.025. The black asterisks in the topography and the shaded areas in the powerspecturm highlight the clusters supporting the rejection of the null hypothesis of no effect between trials dominated by high vs. low gaze variability during encoding. **C-** Rain cloud plots^42^ illustrating the distribution of remembered (left) and forgotten (right) itmes split by gaze variability (high vs. low) during encoding. **D**-The distribution of high (left) and low (right) gaze variability split by memory performance (remembered vs. forgotten).

### Gaze variability and visual cortical activity are continuously coupled

Given the results described above, we next asked whether a co-fluctuation of gaze bias and alpha/beta power would be restricted to the memory task or could be a more general pattern. Specifically, we asked whether alpha/beta power and gaze bias would be correlated over the whole period of the experiment. The results of this analysis are summarized in Figure 3.

**Figure 3:**
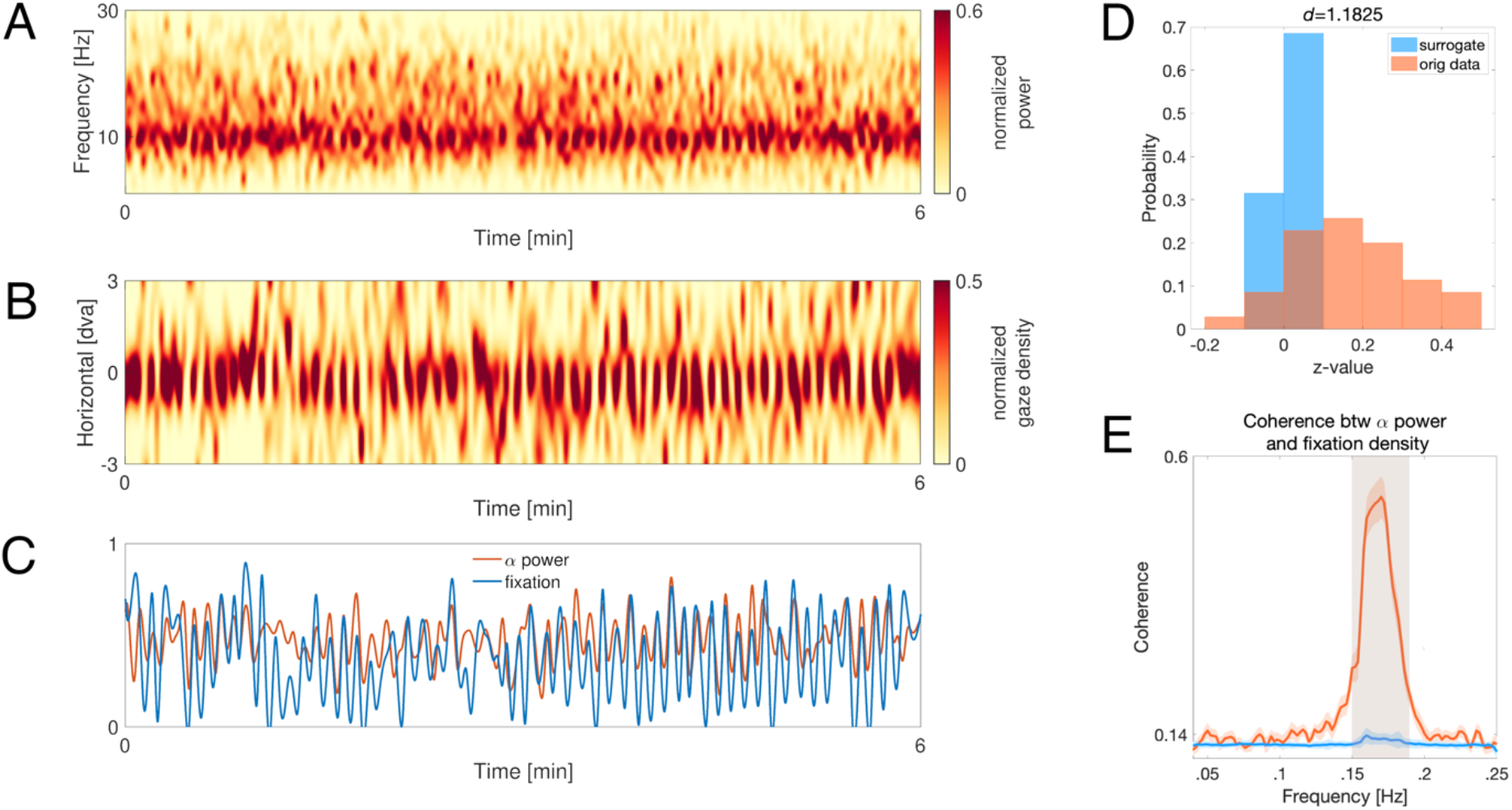
Time course of the covariation of occipital alpha power and gaze direction maintenance. **A-**Single subject example of TFR averaged over occipital sensors, corrected for the presense of the aperiodic 1/F component. X-axis denotes time in minutes and y-axis frequency in Hz. The first 6 minutes of the recording are shown for illustrative purposes. Color code denotes the range corrected power (x – min)/(max-min) varying between 0 and 1. **B**-Gaze density in the horizontal dimension around the fixation location and vertical range of ±3 dva expressed as function of time (x-axis). The middle position of the y-axis denots the fixation location, up rightward gaze bias and down denotes leftward gaze bias. Color code illustrates the range corrected gaze density varying between 0 and 1. **C-** Time course of alpha power (10-14Hz, red color) and gaze density at the fixation location (blue color) overlaid on top of each other. The x-axis is identical to A and B. **D**-Normalized probability histogramms (sum of y-axis equals 1) of the Spearman’s rho correlation coefficient between the time courses of alpha power and gaze density at fixation, converted in z-values. Red color denotes the distribution of all correlations across all participants estimated on the original time series. Blue color denotes the distribution of correlations estimated on circularly shifted time series after 1000 permutations. A significant difference was confirmed t(34)= 6.8, CI[0.12 0.22], p < 7.4e-08 and estimated effect size of Cohen’s d= 1.1541. **E**-Coherence spectra between the alpha power and gaze density timeseries. Frequency is depicted on the x-axis and coherence magnitude on the y-axis. Red color denotes the original time series and the blue color the surrogate data generated after permuting the trial correspondense between alpha power and gaze densitiy data and computing the coherence (1000-fold permutation) and averaging over folds. Shading illsustrates standard error.

First, we computed the time-frequency representation of power (TFR) for the entire recording per participant (M/SEM = 60.59/±2.01 min) and accounted for the presence of aperiodic activity ^43^. For illustrative purposes, Figure 3A illustrates the first 6 minutes in the recording of a representative participant. The TFR depicts the spontaneous power modulation of alpha (predominantly in 10-14 Hz) activity over occipital sensors. In addition, we extracted the fixation density for the horizontal direction averaged across the vertical direction. This fixation density signal can be visualized as a function of time with the horizontal viewing direction now illustrated on the y-axis (Figure 3B). In both panels (Figure 3 A and B), the power of the signals is range corrected (*[x – min]/[max-min]*) to vary between 0 and 1. The time series of alpha/beta power and fixation density can be extracted and overlaid (Figure 3C) conveying a (descriptive) similarity between the variation in alpha/beta power and the variation in fixation maintenance. Using this approach, we computed Spearman’s rho correlations between both timeseries for the entire recording session in each participant. In addition, surrogate data was generated by circularly shifting both time series (1000 times) and computing the corresponding correlation. This analysis yielded two distributions of correlations that can be parametrically compared (Figure 3D). For each participant we derived two correlation coefficients. These correlations obtained from the original data were significantly different from the one generated by the surrogate data (t^(34)^= 6.8, CI[0.12 0.22], p < 7.4e-08, Cohen’s d= 1.1541). This effect was confirmed on the individual participant level, with 33 out of 35 participants showing a significant difference (Figure S3). Finally, we computed the coherence between the two timeseries to infer the frequency of any phase relationship in the covarion of gaze and occipital alpha/beta power. Both time courses were segmented into trials of 200 seconds length, and the coherence across trials between gaze and alpha/beta power time series was computed. We observed a consistent coherence at app. 0.18Hz (Figure 3E, cluster permutation test, p < 0.025, t > 10, Cohen’s d > 1.5) as compared to surrogate data generated after 1000 permutations of the trials. As illustrated in Figure 3E, the permutation procedure effectively eliminated the temporal correspondence between alpha/beta power and gaze density time courses, reflected in a flatter coherence spectrum.

## Discussion

Previous research has shown that gaze patterns as well as amplitude modulation of neural activity in posterior cortex during encoding of visual scenes strongly predicts memory performance. However, gaze patterns and amplitude modulation have thus far been studied in isolation. Our findings indicate that we might be looking at two sides of the same coin: gaze patterns and amplitude modulations covary and jointly predict memory performance. We simultaneously recorded MEG and eye movements in a free viewing memory paradigm. To relate alpha/beta MEG activity as well gaze variability to successful encoding, we contrasted later remembered versus later forgotten study phase items. In line with many previous studies ^18-30^, we find an alpha/beta SME, with greater decreases in 10-20 Hz power for later remembered as compared to later forgotten items. Importantly, we here demonstrate that gaze covaries with this well-established effect. During the presentation of later remembered items, gaze was directed away from the center to a greater extent than during later forgotten items. This gaze bias indicates that more extensive visual exploration correlates with a higher probability to remember the item, a finding in line with previous studies showing that the amount of saccades on complex visual stimuli predicts whether or not the stimulus will be remembered^8^. Just like the alpha/beta SME, the gaze-related SME also correlated with the reported confidence judgement. Items that were later remembered with high confidence displayed stronger gaze biases and stronger alpha/beta desynchronization than those items remembered with lower confidence. Moreover, we find that splitting the encoding EEG data between high vs. low gaze variability trials closely emulates the traditional alpha/beta SME, both in frequency and topography. Trials with high gaze bias display significantly stronger alpha/beta power decreases than trials with lower gaze bias. Contrasting subsequent memory performance based on gaze bias confirmed that higher gaze bias was predictive of successful memory performance. These results indicate that gaze variability and alpha/beta activity go hand in hand in service of memory formation, a notion in line with a previous study linking saccades and the phase of alpha/beta activity successful remembering^44^.

Generalizing the covariation of gaze and alpha/beta activity beyond memory encoding, we show that gaze covaries with spontaneous fluctuations of alpha/beta during the pre-stimulus baseline. Again, greater variation in gaze was correlated with larger decreases in alpha/beta power. Moving even further away from specific, task-related activity, we show that alpha/beta activity and gaze covary over the course of the whole experiment. Including an hour of simultaneous eye tracking and MEG recordings per participant, we show substantial covariation of alpha power and gaze that is task-independent. Taken together, these observations support the idea of a fundamental and domain-general link between visual alpha/beta activity and oculomotor action.

The present results are in line with recent studies indicating a strong link between eye movements and alpha/beta activity. Biases in gaze direction are associated with a topographically consistent decrease of alpha/beta power, independent of the stimulus modality (e.g. vision, audition)^35^, independent of retinal input and present during full darkness^33^. Similar associations between gaze bias and alpha/beta power modulation have recently been reported in the context of working memory^36,45^. Together with the here reported results, these studies make a strong case that the association between gaze and alpha/beta activity reflects a fundamental functional organization linking visual cortical signals to oculomotor action.

The present findings are also in line with previous studies showing a functionally relevant coordination of eye movements and brain activity. Eye movements affect the neural activity in- and outside visual areas^44,46-53^, as well as in brain areas crucial for episodic memory formation: neural activity in the primate hippocampus is sensitive to eye movements^54-58^. In particular, the phase of hippocampal low-frequency activity has been shown to be aligned to saccadic eye movements^59,60^, and the amount of alignment was found to be functionally relevant^61,62^. Such phase alignment could be a general motif in the nervous system to facilitate the organization of neural ensembles^63^. The exact temporal coordination of neural activity and eye movements could be brought about by efference copies^64^. These copies of motor commands branch off corticofugal projections and could, in principle, be distributed across many brain areas. If they are capable of inducing phase alignment, efference copies could trigger the alignment of neural activity across a broad range of areas and, thereby, facilitate neural processing and communication^65-67^.

We here demonstrate a link between eye movements and alpha/beta activity during the free viewing of visual scenes in the study phase of a memory paradigm. Alpha/beta SMEs during memory encoding are indeed a common finding, yet to what extent the present linkage between alpha/beta SME and gaze variability generalize across stimulus material such as faces and words^68^ vs. visual scenes requires further examination. Moreover, one could argue that the present findings are limited to free viewing encoding conditions, as the majority of the studies investigating SMEs ask their research participants to maintain fixation. Hence, the observation of alpha/beta variation with gaze variability might be somewhat coincidental. We hypothesize that the present findings will generalize to such conditions as well. This prediction is based on the present observation that the alpha/beta-gaze relationship was, albeit weaker, also evident when the eyes do not move feely (e.g. maintain fixation during the baseline period, Figure 1E,F). As fixation is not a stationary event but a process involving the control of miniature eye movements such as micro saccades^69^, a testable prediction is that independent of task instructions (fixation, free view), alpha/beta SME modulation should be also associated with gaze-related SME. An empirical question that can be examined in future studies.

In conclusion, we demonstrate that alpha/beta activity (10-20 Hz) in visual cortex and eye movements covary and jointly predict subsequent memory. Going beyond memory-related brain activity, we show that this covariation is task-independent and preserved over the course of hours. The present results thus support a complementary view on the role of alpha/beta activity, emphasizing its fundamental interrelation with oculomotor behavior.

## Acknowledgements

The study was performed at the Donders Institute for Brain, Cognition and Behaviour. The authors would like to thank Ole Jensen and Christian F. Doeller for their involvement in conceptualizing the study. The authors would also like to thank all the participants volunteering in the experiment.

## Materials and Methods

Part of the present dataset has been previously analyzed and documented elsewhere^44^.

### Ethic statement

Study enrollment followed the approval of the local ethics committee-commission for human related research CMO-2014/288 region Arnhem/Nijmegen, the Netherlands. Prior to participation all volunteers were given written informed consent in accordance with the Declaration of Helsinki.

### Participants

A total of 48 participants were recruited. Thirty-five were included in the present study. Thirteen participants were excluded due to: not completing the study (N=7), excessive artifacts (N=3) and technical difficulties during acquisition (N=3). The included participants sample (24 female, age range 18-30, mean age 23.1y) reported no history of neurological and/or psychiatric diagnosis and had normal or corrected to normal vision.

### Task design and procedure

Visual scenes were presented to the participants projected onto a back-projection screen position in front of the participants inside a magnetically shielding room (MSR). The back-projection screen had the size of 39 × 46 cm corresponding to a visual angle of vertical 27° × 32° horizontal dva (degrees of visual angle). Three sets of visual scenes (100 scenes each) were used. Two sets were displayed during the study (encoding) and the test phase. The scenes in the third set served as lures and were presented exclusively during the test phase. Set assignment to study and test phase was counterbalanced across participants. A training session preceded the actual memory task to ensure participants familiarity with the task procedures. Nine additional scenes not used in the main experiment were shown during the training session. All participants were made aware about the memory test prior to data acquisition.

During the study phase, visual scenes (indoor and outdoor images) were displayed for 4 sec with the constrained that no more than 4 scenes from the same category could appear consecutively. Participants were instructed to to report via button press whether or not the current scene displayed was an indoor or outdoor image. The response was given during the display of a fixation cross with a variable duration between 1-2 sec presented after each scene. During scene viewing, participants were not required to maintain fixation but were allowed to freely explore the scenes.

A short distracter session was required after each study phase. This session consisted of solving simple mathematical problems (appr. 1 min), a simple saccade task during which participants had to saccade to various locations on the screen (5 min), eyes open and eyes closed data acquisition (appr. 1 min each). The purpose of this distracter session was to prevent participants from covert rehearsing.

The test phase followed thereafter. The three sets of images (200 old items from the study phase intermixed with 100 new items) were randomly presented for viewing duration of 4 sec. The randomization had the constrained that no more than 4 images of the same type (old/new) could be shown consecutively. Following each image presentation, a 6-point response scale was displayed. Participants were required to indicate whether the just presented image was “old” or “new” with the scale ranging from “very sure old”(1) to “very sure new”(6). The 6-point scale remained on the visual display until the participants response was given followed by a fixation cross of variable duration ranging between 0.75-1.25 sec.

### Data acquisition

Whole head magnetoencephalography (MEG) was acquired with a 275-axial gradiometer system (VSM MedTech/CTF MEG, Coquitlam, Canada) within the MSR. During data acquisition, the sampling frequency was set to 1200 Hz using a low-pass antialiasing filter at a cutoff frequency of 300 Hz. Ocular artifacts were monitored by horizontal and vertical electrooculogram using Ag/AgCl electrodes and a bipolar montage. Head movements were tracked continuously using 3 coils placed in the vicinity of the nasion and the left and right ear canals^70^. An Eyelink 1000 (SR Research) system was used to monitor horizontal and vertical eye movements of the left eye. Prior to data acquisition, eye tracker calibration was performed which involved the collection of gaze fixation samples from pre-defined positions (9 dots on a 3×3 grid) on the visual display. Raw eye tracking data was mapped to these pre-defined screen coordinates followed by a validation procedure to ensure sufficient correspondence between the position of the current gaze fixations and the one obtained during the preceding calibration. The calibration was accepted if the difference was < 1° dva.

### Data preprocessing and analysis

#### MEG data

Offline data analyses was performed with the open source software for neuroelectric- and neuromagnetic data analysis (FieldTrip^71^). The continuous data was segmented around the events of interest (scene onsets) into epochs of 8 sec (3 sec baseline prior to event onset). Following a finite impulse response band-pass filter (1-40Hz) the epochs were re-segmented to exclude potential filter artifacts resulting in time range of -2.5 to 4.5 sec around the event onset. Oculo-muscular and cardiac artifacts were identified and removed from further analysis by means of independent component analysis (ICA). These procedures were applied to all data epochs extracted during the study (encoding) and test (retrieval) phase. Subsequently, for each epoch, time-frequency estimates of power were computed using a sliding window of 0.5 sec and a Hanning taper resulting in a frequency resolution of appr. 2Hz. The window slid every 50 ms within the range of -2 to 4 sec around the event onset. Power estimates were averaged across epochs for each condition separately, e.g. encoding (remembered/forgotten), retrieval (remembered/forgotten). Baseline correction using the time window of -1 to -.25 (encoding) and -.75 to -.25 (retrieval) was applied transforming the raw power estimates into dB change from pre-stimulus baseline. The number of trials used in the present analysis were N^remembered^ = 145.15/30 (M/STD) and N^forgotten^ = 54.77/30 (M/STD).

We followed a similar approach to split the epochs into high versus low pre-stimulus alpha power during scene encoding. Specifically, a Fast Fourier transform was applied to the data prior to stimulus onset (-1 to 0 sec), including all occipital sensors (see Figure S4). A Hanning taper was used, resulting in appr. 1 Hz frequency resolution. The mean power within the alpha/beta frequency range (10-20Hz) was extracted and averaged over all occipital sensors. Subsequently, trials with high and low alpha power were separated by means of median split. Using this subset of high and low alpha power trials, the power estimates were re-grouped and averaged over trials. Finally, the difference between the condition’s alpha power low minus alpha power high was computed.

#### Gaze data

The raw eye tracking data was converted form voltage to pixel coordinates following the procedures described here https://www.fieldtriptoolbox.org/getting_started/eyelink/#whatare-the-units-of-the-eye-tracker-data. Gaze density was expressed as the 2D heat map according to the procedures described here (https://stackoverflow.com/questions/46996206/matlab-creating-a-heatmap-to-visualize-density-of-2d-point-data). Briefly, the scatter plot of all *x* and *y* positions was converted into an image array (using *imagesc*.*m* in MATLAB) after a 2-D convolution (*conv2*.*m*) with a Gaussian filter matrix *G*. For the given range of *x* and *y* coordinates denoted as *xG* and *yG* and width parameter sigma (in the present case set to 2.5), *G* = exp[-xG^2^/(2*sigma^2^) – yG^2^/(2*sigma^2^)]. After 2D conversion the eye tracking data was converted to a structure that can be read by FieldTrip akin to the one for time-frequency data. Subsequently, all approaches available for statistical treatment of time-frequency data could be applied to the gaze density data.

#### Relationships between MEG and gaze data

In order to examine the relationship between the continuous MEG recording and the gaze variation as presented in Figure 2 the following procedures were utilized. First, the continuous data form occipital sensors was segmented into trials of 2 sec length. Following Fourier analysis as described above, the 1/f aperiodic component was removed from each trial using the *specparam* routines^72^ allowing the parametrization and visualization of periodic components (e.g. alpha activity) in the continuous data. Subsequently, these trials were concatenated yielding a time-frequency representation of power (e.g. Figure 2A). Next, for each trial, the corresponding gaze data was computed as described above, averaged along the vertical direction and concatenated, yielding a time by horizontal position spectrogram with color coded density of horizontal gaze direction (e.g. Figure 2B). For each participant, the time course of occipital alpha power was extracted by averaging the spectral estimates across the dominant frequency band width (e.g. 10-14Hz) across all occipital sensors. The reason for taking 10-14Hz rather than alpha/beta activity is that in spontaneous recordings the dominant rhythm is alpha, whereas alpha/beta decreases are scored as function of baseline or some contrast that in this case we do not have. Instead we took the spontaneous signal exhibiting strongest power mostly in the 10-14Hz range. Similarly, the time course of gaze density averaged around fixation at 0 dva (± .5 horizontal dva) was extracted. These time courses were correlated using a bootstrapping resampling procedure with 1000 iterations yielding a distribution of correlations between the two time courses (e.g. Figure 2D). Surrogate data was generated by circularly shifting the time series 1000 times, where at each iteration, the length of the time shift was randomly chosen ranging between 1s and the total duration of the recording. This procedure resulted in a surrogate distribution of correlations (e.g. Figure 2D), against which the distribution of correlations obtained from the original data was compared. All correlation coefficients were transformed into z-scores (e.g. *z = log((1+r) / (1-r))/2*) prior to further statistical evaluation. Finally, the coherence between the MEG and gaze time series was computed. Given a sampling frequency of 0.5 Hz determined by the length of the trials used to derived these time courses (see above), the MEG and gaze data was first re-segmented into epochs of 200s length. A multi-taper approach^73^ was used to estimate power and cross-spectral density for each trial padded with zeros at the begin and the end of each trial (500 sec in total). Three tapers were used covering the frequency range from 0 to Nyquist frequency (0.25Hz). The phase in the cross-spectra represents the phase difference between the oscillatory signals of the MEG and gaze time series, with consistent phase difference resulting in larger coherence values. Surrogate data was generated by permuting the trial order of the MEG and gaze data with respect to each other and computing coherence. This approach was repeated 1000 times and the resulting coherence values were averaged resulting in the surrogate coherence result presented in Figure 2E.

#### Statistics

Statistical control followed the cluster-based permutation framework ^39^. This approach utilizes clustering across sensors, time points, and frequency (wherever appropriate). Multiple comparisons problem was addressed by using 1000 permutations and a two-tailed alpha threshold of 0.05 (p < 0.025). Correlations were evaluated by utilizing Spearman’s correlation coefficient Rho.

## References

1 Otero-Millan, J., Troncoso, X. G., Macknik, S. L., Serrano-Pedraza, I. & Martinez-Conde, S. Saccades and microsaccades during visual fixation, exploration, and search: foundations for a common saccadic generator. J Vis 8, 21 21–18, doi:10.1167/8.14.21 (2008).

2 Damiano, C. & Walther, D. B. Distinct roles of eye movements during memory encoding and retrieval. Cognition 184, 119–129, doi:10.1016/j.cognition.2018.12.014 (2019).

3 Fehlmann, B. et al. Visual Exploration at Higher Fixation Frequency Increases Subsequent Memory Recall. Cerebral cortex communications 1, tgaa032, doi:10.1093/texcom/tgaa032 (2020).

4 Molitor, R. J., Ko, P. C., Hussey, E. P. & Ally, B. A. Memory-related eye movements challenge behavioral measures of pattern completion and pattern separation. Hippocampus 24, 666–672, doi:10.1002/hipo.22256 (2014).

5 Olsen, R. K. et al. The relationship between eye movements and subsequent recognition: Evidence from individual differences and amnesia. Cortex; a journal devoted to the study of the nervous system and behavior 85, 182–193, doi:10.1016/j.cortex.2016.10.007 (2016).

6 Kragel, J. E. & Voss, J. L. Looking for the neural basis of memory. Trends in cognitive sciences 26, 53–65, doi:10.1016/j.tics.2021.10.010 (2022).

7 Broers, N., Bainbridge, W. A., Michel, R., Balestrieri, E. & Busch, N. A. The extent and specificity of visual exploration determines the formation of recollected memories in complex scenes. J Vis 22, 9, doi:10.1167/jov.22.11.9 (2022).

8 Voss, J. L., Bridge, D. J., Cohen, N. J. & Walker, J. A. A Closer Look at the Hippocampus and Memory. Trends in cognitive sciences 21, 577–588, doi:10.1016/j.tics.2017.05.008 (2017).

9 Johansson, R., Nystrom, M., Dewhurst, R. & Johansson, M. Eye-movement replay supports episodic remembering. Proc Biol Sci 289, 20220964, doi:10.1098/rspb.2022.0964 (2022).

10 Wynn, J. S., Ryan, J. D. & Buchsbaum, B. R. Eye movements support behavioral pattern completion. Proc Natl Acad Sci U S A 117, 6246–6254, doi:10.1073/pnas.1917586117 (2020).

11 Wynn, J. S., Shen, K. & Ryan, J. D. Eye Movements Actively Reinstate Spatiotemporal Mnemonic Content. Vision (Basel) 3, doi:10.3390/vision3020021 (2019).

12 Brandt, S. A. & Stark, L. W. Spontaneous eye movements during visual imagery reflect the content of the visual scene. Journal of cognitive neuroscience 9, 27–38, doi:10.1162/jocn.1997.9.1.27 (1997).

13 Bochynska, A. & Laeng, B. Tracking down the path of memory: eye scanpaths facilitate retrieval of visuospatial information. Cogn Process 16 Suppl 1, 159–163, doi:10.1007/s10339-015-0690-0 (2015).

14 Laeng, B. & Teodorescu, D.-S. Eye scanpaths during visual imagery reenact those of perception of the same visual scene. Cognitive science 26, 207–231, doi:https://doi.org/10.1207/s15516709cog2602_3 (2002).

15 Gbadamosi, J. & Zangemeister, W. H. Visual imagery in hemianopic patients. Journal of cognitive neuroscience 13, 855–866, doi:10.1162/089892901753165782 (2001).

16 Johansson, R., Holsanova, J. & Holmqvist, K. Pictures and spoken descriptions elicit similar eye movements during mental imagery, both in light and in complete darkness. Cognitive science 30, 1053–1079, doi:10.1207/s15516709cog0000_86 (2006).

17 Paller, K. A. & Wagner, A. D. Observing the transformation of experience into memory. Trends in cognitive sciences 6, 93–102, doi:10.1016/s1364-6613(00)01845-3 (2002).

18 Vogelsang, D. A., Gruber, M., Bergström, Z. M., Ranganath, C. & Simons, J. S. Alpha Oscillations during Incidental Encoding Predict Subsequent Memory for New “Foil” Information. Journal of cognitive neuroscience 30, 667–679, doi:10.1162/jocn_a_01234 (2018).

19 Fellner, M. C., Bäuml, K. H. & Hanslmayr, S. Brain oscillatory subsequent memory effects differ in power and long-range synchronization between semantic and survival processing. NeuroImage 79, 361–370, doi:10.1016/j.neuroimage.2013.04.121 (2013).

20 Khader, P. H., Jost, K., Ranganath, C. & Rosler, F. Theta and alpha oscillations during working-memory maintenance predict successful long-term memory encoding. Neurosci Lett 468, 339–343, doi:10.1016/j.neulet.2009.11.028 (2010).

21 Hanslmayr, S., Spitzer, B. & Bäuml, K.-H. Brain Oscillations Dissociate between Semantic and Nonsemantic Encoding of Episodic Memories. Cerebral Cortex 19, 1631–1640, doi:10.1093/cercor/bhn197 (2008).

22 Klimesch, W. et al. Event-related desynchronization (ERD) and the Dm effect: does alpha desynchronization during encoding predict later recall performance? Int J Psychophysiol 24, 47–60, doi:10.1016/s0167-8760(96)00054-2 (1996).

23 Sederberg, P. B., Kahana, M. J., Howard, M. W., Donner, E. J. & Madsen, J. R. Theta and gamma oscillations during encoding predict subsequent recall. J Neurosci 23, 10809–10814 (2003).

24 Hanslmayr, S., Staudigl, T. & Fellner, M. C. Oscillatory power decreases and long-term memory: the information via desynchronization hypothesis. Front Hum Neurosci 6, 74, doi:10.3389/fnhum.2012.00074 (2012).

25 Griffiths, B. J. et al. Alpha/beta power decreases track the fidelity of stimulus-specific information. Elife 8, doi:10.7554/eLife.49562 (2019).

26 Griffiths, B. J., Martín-Buro, M. C., Staresina, B. P. & Hanslmayr, S. Disentangling neocortical alpha/beta and hippocampal theta/gamma oscillations in human episodic memory formation. NeuroImage 242, 118454, doi:https://doi.org/10.1016/j.neuroimage.2021.118454 (2021).

27 Strunk, J. & Duarte, A. Prestimulus and poststimulus oscillatory activity predicts successful episodic encoding for both young and older adults. Neurobiology of aging 77, 1–12, doi:10.1016/j.neurobiolaging.2019.01.005 (2019).

28 Sander, M. C., Fandakova, Y., Grandy, T. H., Shing, Y. L. & Werkle-Bergner, M. Oscillatory Mechanisms of Successful Memory Formation in Younger and Older Adults Are Related to Structural Integrity. Cerebral Cortex 30, 3744–3758, doi:10.1093/cercor/bhz339 (2020).

29 Griffiths, B. J., Martín-Buro, M. C., Staresina, B. P., Hanslmayr, S. & Staudigl, T. Alpha/beta power decreases during episodic memory formation predict the magnitude of alpha/beta power decreases during subsequent retrieval. Neuropsychologia 153, 107755, doi:10.1016/j.neuropsychologia.2021.107755 (2021).

30 Hanslmayr, S. & Staudigl, T. How brain oscillations form memories--a processing based perspective on oscillatory subsequent memory effects. NeuroImage 85 Pt 2, 648–655, doi:10.1016/j.neuroimage.2013.05.121 (2014).

31 Parish, G., Hanslmayr, S. & Bowman, H. The Sync/deSync Model: How a Synchronized Hippocampus and a Desynchronized Neocortex Code Memories. The Journal of Neuroscience 38, 3428–3440, doi:10.1523/jneurosci.2561-17.2018 (2018).

32 Hanslmayr, S., Staresina, B. P. & Bowman, H. Oscillations and Episodic Memory: Addressing the Synchronization/Desynchronization Conundrum. Trends in neurosciences 39, 16–25, doi:10.1016/j.tins.2015.11.004 (2016).

33 Popov, T., Miller, G. A., Rockstroh, B., Jensen, O. & Langer, N. Alpha oscillations link action to cognition: An oculomotor account of the brain’s dominant rhythm. bioRxiv, 2021.2009.2024.461634, doi:10.1101/2021.09.24.461634 (2021).

34 Staudigl, T., Hartl, E., Noachtar, S., Doeller, C. F. & Jensen, O. Saccades are phase-locked to alpha oscillations in the occipital and medial temporal lobe during successful memory encoding. PLoS biology 15, e2003404 (2017).

35 Popov, T., Langer, N., Gips, B., Weisz, N. & Jensen, O. Sound-location specific alpha power modulation in the visual cortex in absence of visual input. bioRxiv, 2021.2003.2015.435371, doi:10.1101/2021.03.15.435371 (2021).

36 Printzlau, F. A. B., Myers, N. E., Manohar, S. G. & Stokes, M. G. Neural Reinstatement Tracks Spread of Attention between Object Features in Working Memory. Journal of cognitive neuroscience, 1–21, doi:10.1162/jocn_a_01879 (2022).

37 Popov, T., Gips, B., Weisz, N. & Jensen, O. Brain areas associated with visual spatial attention display topographic organization during auditory spatial attention. Cerebral cortex (New York, N.Y. : 1991), doi:10.1093/cercor/bhac285 (2022).

38 Liu, B., Nobre, A. C. & van Ede, F. Microsaccades transiently lateralise EEG alpha activity. bioRxiv, 2022.2009.2002.506318, doi:10.1101/2022.09.02.506318 (2022).

39 Maris, E. & Oostenveld, R. Nonparametric statistical testing of EEG- and MEG-data. J Neurosci Methods 164, 177–190, doi:10.1016/j.jneumeth.2007.03.024 (2007).

40 Dimigen, O., Sommer, W., Hohlfeld, A., Jacobs, A. M. & Kliegl, R. Coregistration of eye movements and EEG in natural reading: analyses and review. J Exp Psychol Gen 140, 552–572, doi:10.1037/a0023885 (2011).

41 Engbert, R. & Mergenthaler, K. Microsaccades are triggered by low retinal image slip. Proc Natl Acad Sci U S A 103, 7192–7197, doi:10.1073/pnas.0509557103 (2006).

42 Allen, M., Poggiali, D., Whitaker, K., Marshall, T. R. & Kievit, R. A. Raincloud plots: a multi-platform tool for robust data visualization. Wellcome Open Res 4, 63, doi:10.12688/wellcomeopenres.15191.1 (2019).

43 Donoghue, T. et al. Parameterizing neural power spectra into periodic and aperiodic components. Nature Neuroscience 23, 1655–1665, doi:10.1038/s41593-020-00744-x (2020).

44 Staudigl, T., Hartl, E., Noachtar, S., Doeller, C. F. & Jensen, O. Saccades are phase-locked to alpha oscillations in the occipital and medial temporal lobe during successful memory encoding. PLoS Biol 15, e2003404, doi:10.1371/journal.pbio.2003404 (2017).

45 Liu, B., Nobre, A. C. & van Ede, F. Functional but not obligatory link between microsaccades and neural modulation by covert spatial attention. Nature Communications 13, 3503, doi:10.1038/s41467-022-31217-3 (2022).

46 Brunet, N. et al. Visual cortical gamma-band activity during free viewing of natural images. Cerebral cortex (New York, N.Y. : 1991) 25, 918–926, doi:10.1093/cercor/bht280 (2015).

47 McFarland, J. M., Bondy, A. G., Saunders, R. C., Cumming, B. G. & Butts, D. A. Saccadic modulation of stimulus processing in primary visual cortex. Nature Communications 6, 8110, doi:10.1038/ncomms9110 (2015).

48 Ito, J., Maldonado, P., Singer, W. & Grün, S. Saccade-related modulations of neuronal excitability support synchrony of visually elicited spikes. Cerebral cortex (New York, N.Y. : 1991) 21, 2482–2497, doi:10.1093/cercor/bhr020 (2011).

49 Rajkai, C. et al. Transient cortical excitation at the onset of visual fixation. Cerebral cortex (New York, N.Y. : 1991) 18, 200–209, doi:10.1093/cercor/bhm046 (2008).

50 Barczak, A. et al. Dynamic Modulation of Cortical Excitability during Visual Active Sensing. Cell reports 27, 3447-3459.e3443, doi:10.1016/j.celrep.2019.05.072 (2019).

51 Leszczynski, M. et al. Neural activity in the human anterior thalamus during natural vision. Scientific Reports 11, 17480, doi:10.1038/s41598-021-96588-x (2021).

52 Hamamé, C. M. et al. Functional selectivity in the human occipitotemporal cortex during natural vision: evidence from combined intracranial EEG and eye-tracking. NeuroImage 95, 276–286, doi:10.1016/j.neuroimage.2014.03.025 (2014).

53 Purpura, K. P., Kalik, S. F. & Schiff, N. D. Analysis of perisaccadic field potentials in the occipitotemporal pathway during active vision. Journal of neurophysiology 90, 3455–3478, doi:10.1152/jn.00011.2003 (2003).

54 Andrillon, T., Nir, Y., Cirelli, C., Tononi, G. & Fried, I. Single-neuron activity and eye movements during human REM sleep and awake vision. Nature Communications 6, 7884, doi:10.1038/ncomms8884 (2015).

55 Mao, D. et al. Spatial modulation of hippocampal activity in freely moving macaques. Neuron 109, 3521-3534.e3526, doi:https://doi.org/10.1016/j.neuron.2021.09.032 (2021).

56 Katz, C. N. et al. A corollary discharge mediates saccade-related inhibition of single units in mnemonic structures of the human brain. Curr Biol 32, 3082-3094.e3084, doi:10.1016/j.cub.2022.06.015 (2022).

57 Wagner, I. C., Jensen, O., Doeller, C. F. & Staudigl, T. Saccades are coordinated with directed circuit dynamics and stable but distinct hippocampal patterns that promote memory formation. bioRxiv, 2022.2008.2018.504386, doi:10.1101/2022.08.18.504386 (2022).

58 Liu, Z. X., Shen, K., Olsen, R. K. & Ryan, J. D. Visual Sampling Predicts Hippocampal Activity. J Neurosci 37, 599–609, doi:10.1523/JNEUROSCI.2610-16.2016 (2017).

59 Hoffman, K. L. et al. Saccades during visual exploration align hippocampal 3-8 Hz rhythms in human and non-human primates. Frontiers in systems neuroscience 7, 43, doi:10.3389/fnsys.2013.00043 (2013).

60 Doucet, G., Gulli, R. A., Corrigan, B. W., Duong, L. R. & Martinez-Trujillo, J. C. Modulation of local field potentials and neuronal activity in primate hippocampus during saccades. Hippocampus 30, 192–209, doi:10.1002/hipo.23140 (2020).

61 Jutras, M. J., Fries, P. & Buffalo, E. A. Oscillatory activity in the monkey hippocampus during visual exploration and memory formation. Proc Natl Acad Sci U S A 110, 13144–13149, doi:10.1073/pnas.1302351110 (2013).

62 Staudigl, T., Minxha, J., Mamelak, A. N., Gothard, K. M. & Rutishauser, U. Saccade-related neural communication in the human medial temporal lobe is modulated by the social relevance of stimuli. Science advances 8, eabl6037, doi:10.1126/sciadv.abl6037 (2022).

63 Voloh, B. & Womelsdorf, T. A Role of Phase-Resetting in Coordinating Large Scale Neural Networks During Attention and Goal-Directed Behavior. Frontiers in systems neuroscience 10, 18, doi:10.3389/fnsys.2016.00018 (2016).

64 Sommer, M. A. & Wurtz, R. H. A pathway in primate brain for internal monitoring of movements. Science (New York, N.Y.) 296, 1480–1482, doi:10.1126/science.1069590 (2002).

65 Buzsáki, G. Neural syntax: cell assemblies, synapsembles, and readers. Neuron 68, 362–385, doi:10.1016/j.neuron.2010.09.023 (2010).

66 Fries, P. Rhythms for Cognition: Communication through Coherence. Neuron 88, 220–235, doi:10.1016/j.neuron.2015.09.034 (2015).

67 Schneider, M. et al. A mechanism for inter-areal coherence through communication based on connectivity and oscillatory power. Neuron 109, 4050-4067.e4012, doi:10.1016/j.neuron.2021.09.037 (2021).

68 Fellner, M. C. et al. Spectral fingerprints or spectral tilt? Evidence for distinct oscillatory signatures of memory formation. PLoS Biol 17, e3000403, doi:10.1371/journal.pbio.3000403 (2019).

69 Martinez-Conde, S., Otero-Millan, J. & Macknik, S. L. The impact of microsaccades on vision: towards a unified theory of saccadic function. Nature reviews. Neuroscience 14, 83–96, doi:10.1038/nrn3405 (2013).

70 Stolk, A., Todorovic, A., Schoffelen, J. M. & Oostenveld, R. Online and offline tools for head movement compensation in MEG. NeuroImage 68, 39–48, doi:10.1016/j.neuroimage.2012.11.047 (2013).

71 Oostenveld, R., Fries, P., Maris, E. & Schoffelen, J. M. FieldTrip: Open source software for advanced analysis of MEG, EEG, and invasive electrophysiological data. Comput Intell Neurosci 2011, 156869, doi:10.1155/2011/156869 (2011).

72 Donoghue, T. et al. Parameterizing neural power spectra into periodic and aperiodic components. Nat Neurosci 23, 1655–1665, doi:10.1038/s41593-020-00744-x (2020).

73 Mitra, P. P. & Pesaran, B. Analysis of Dynamic Brain Imaging Data. Biophysical Journal 76, 691–708, doi:https://doi.org/10.1016/S0006-3495(99)77236-X (1999).

