## Supplemental material for "Cortico-ocular coupling in the service of episodic memory formation"

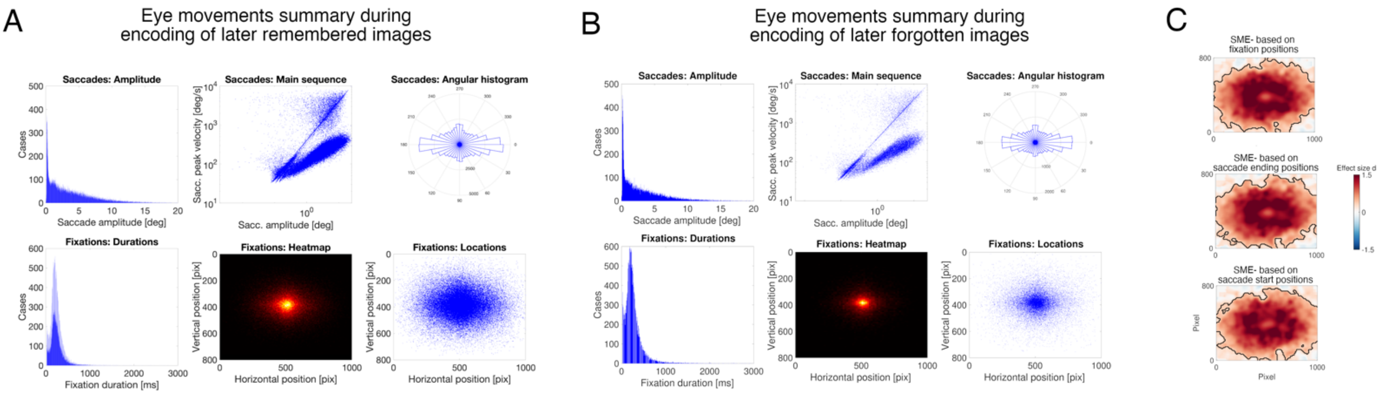


Figure S1: Description of eye movement events across all participants during A- later remembered, B- later forgotten images and C the corresponding gaze bias heat maps.

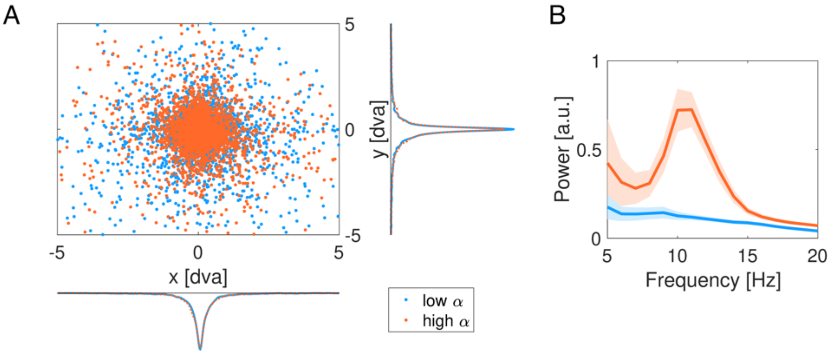


Figure S2: A- distribution of fixations for low (blue) and high (orange) alpha activity. B- power spectra over occipital sensors for high (orange) and low (blue) alpha activity.

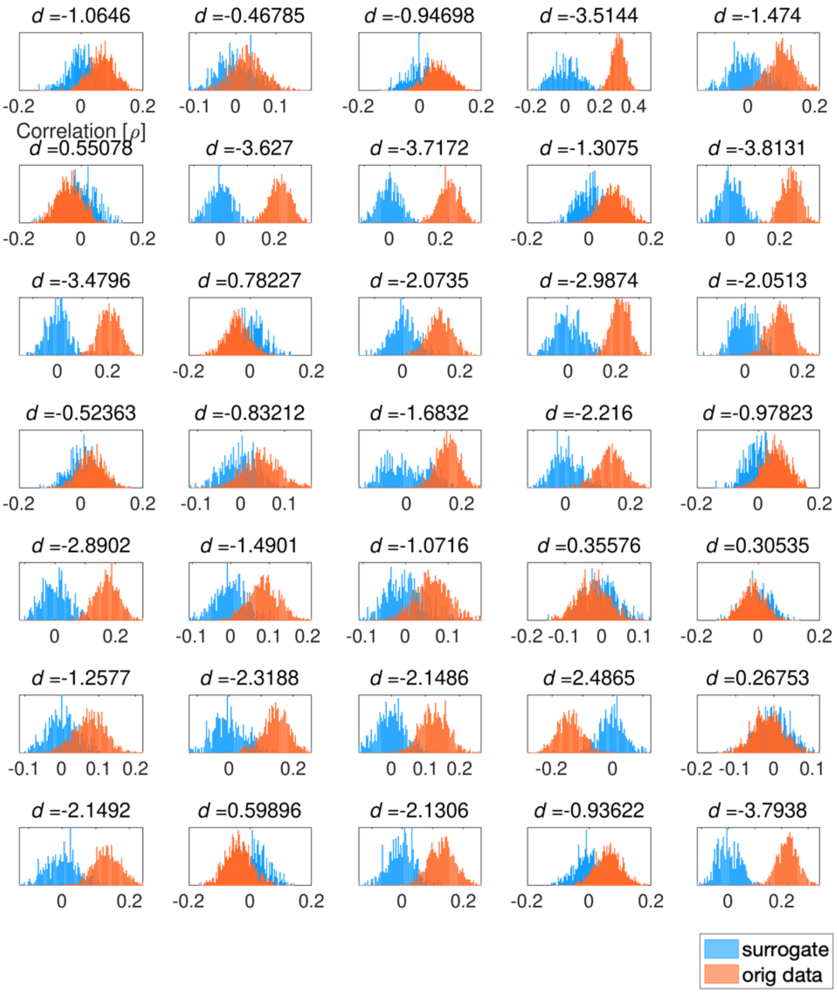


Figure S3: Single participant's distributions of the correlations between gaze and alpha time courses.

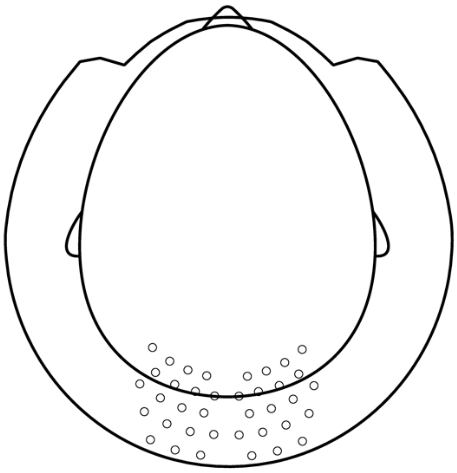

Figure S4: Position of occipital sensors.
